# Glucocorticoid receptor condensates link DNA-dependent receptor dimerization and transcriptional transactivation

**DOI:** 10.1101/2020.11.10.376327

**Authors:** Filipp Frank, Xu Liu, Eric A. Ortlund

## Abstract

The glucocorticoid receptor (GR) is a ligand-regulated transcription factor (TF) that controls the tissue- and gene-specific transactivation and transrepression of thousands of target genes. Distinct GR DNA binding sequences with activating or repressive activities have been identified, but how they modulate transcription in opposite ways is not known. We show that GR forms phase-separated condensates that specifically concentrate known co-regulators via their intrinsically disordered regions (IDRs) *in vitro.* A combination of dynamic, multivalent (between IDRs) and specific, stable interactions (between LxxLL motifs and the GR ligand binding domain) control the degree of recruitment. Importantly, GR DNA-binding directs the selective partitioning of co-regulators within GR condensates such that activating DNAs cause enhanced recruitment of co-activators. Our work shows that condensation controls GR function by modulating co-regulator recruitment and provides a mechanism for the up- and down-regulation of GR target genes controlled by distinct DNA recognition elements.

## Introduction

The glucocorticoid receptor (GR) is a ligand-regulated vertebrate transcription factor that inhibits inflammation and regulates the body’s stress response and metabolism. Synthetic glucocorticoids are used to treat various conditions including autoimmune disorders, allergies and asthma, adrenal insufficiency, heart failure, cancer, and skin conditions. Long-term use, however, causes severe side effects, limiting the use of GCs in chronic conditions.

GR is a multivalent protein with a modular domain architecture containing a combination of intrinsically disordered and stable, folded domains, which is characteristic of the nuclear receptor (NR) family (Figure 1A). The disordered N-terminal domain contains an autonomous activation function (AF1) that mediates interactions with co-regulators. The DNA binding domain (DBD) targets NRs to specific genomic loci. It is connected via a short, intrinsically disordered hinge region to the ligand binding domain (LBD), which contains a second activation function (AF2) responsible for the ligand-dependent recruitment of transcriptional co-regulators(Lonard and O’Malley, 2012; Millard et al., 2013). Co-regulators are broadly classified as co-repressors and co-activators depending on their effects on transcription. They associate with the AF2 via short peptide motifs usually found in their IDRs – the NR box (LxxLL) in co-activators and the CoRNR box (LxxH/IIxxxI/L) in co-repressors. Hundreds of co-regulators have been identified, many of which utilize this same binding surface on the NR LBD. The recruitment of co-regulators to genomic loci by NRs is highly context-dependent so that distinct co-activator and co-repressor complexes are assembled in a cell-type- and gene-specific manner. Little is known, however, about how selectivity in co-regulator recruitment is achieved hampering our ability to predict the magnitude or direction of expression at a given promoter.

**Figure 1:**
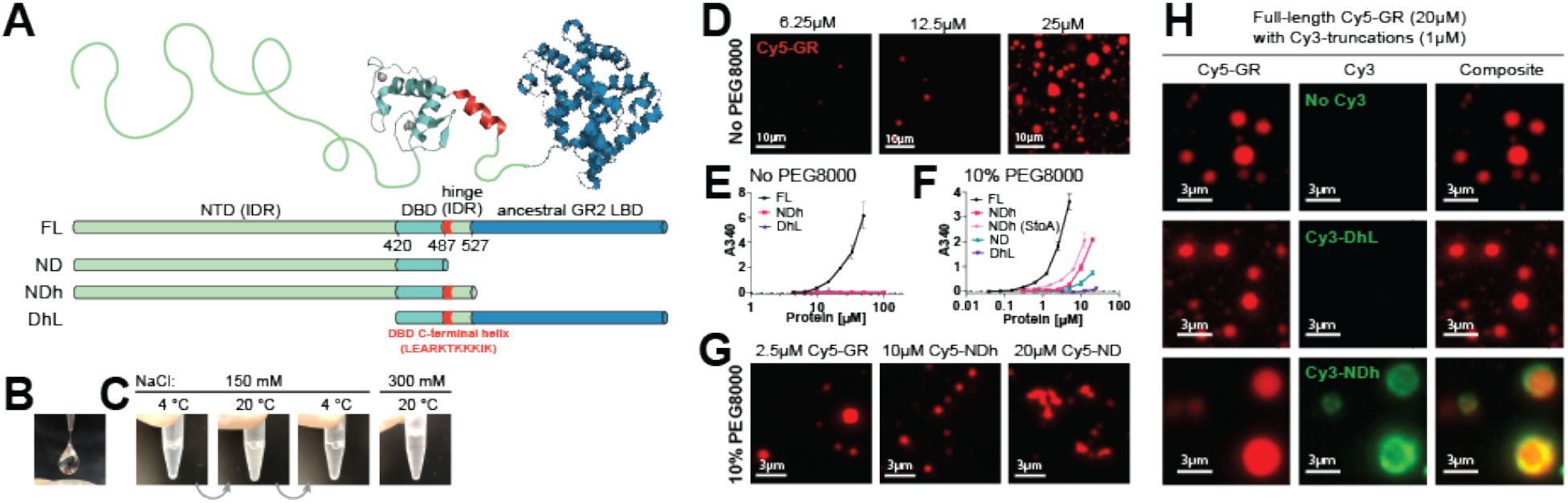
GR forms phase separated condensates in vitro. **(A)** Domain structure of GR and the different recombinant protein constructs used in this study. **(B)** At concentrations above ~1μg/μl GR forms strings when extruded from a pipette tip. **(C)** GR solutions turn turbid at room temperature and reversibly clear up when placed back on ice. Increasing the salt concentration prevents turbidity. **(D)** GR forms well-defined spherical droplets at increasing concentrations. **(E)** Quantification of turbidity in solutions of increasing full-length GR concentrations. Solutions of truncation constructs are not turbid, even at high concentrations. **(F)** A crowding reagent (10% PEG8000) reduces the critical concentration for phase separation in full-length GR and induces turbidity in truncation constructs containing the NTD. **(G)** In the presence of PEG8000 full-length GR and NDh form well-defined spherical condensates, whereas ND forms aggregates. **(H)** NDh is recruited to the surface of GR condensates. A construct missing the NTD is not recruited into GR condensates. *Statistics: Graphs in panels (E) and (F) represent the +/− s.e.m. of at least 4 measurements.*

GR interacts with multiple different types of DNA binding sites *in vivo(Weikum et al., 2017a).* The mode of interaction modulates the transcriptional outcome, i.e. transactivation or transrepression. Canonical glucocorticoid response elements (GREs) are pseudo-palindromic sequences containing two copies of the AGAACA hexamer separated by a three base-pair spacer and generally induce transactivation. GR binds to these sequences as a head-to-head dimer (Figure 3A) in which interprotein contacts provide binding cooperativity (Luisi et al., 1991; Meijsing et al., 2009; Watson et al., 2013). The more recently discovered negative glucocorticoid response element contains an inverted repeat separated by a short spacer of up to 2 base-pairs (CTCC(N)0-2GGAGA) and causes transrepression(Surjit et al., 2011). Two GR molecules bind to this sequence on opposite sides of the DNA without contacts between the DBDs and with negative cooperativity(Hudson et al., 2013). A third class of GR recognition sequences contain canonical half-sites (AGAACA), which GR engages as a monomer(Lim et al., 2015; Schiller et al., 2014). Similarly, GR binds to cryptic half-sites (AATTY, with Y representing a pyrimidine base) found in genomic NF-κB response elements (κBREs), which drives transrepression of many inflammatory genes(Hudson et al., 2018b). Finally, GR can repress genes without directly contacting DNA by “tethering” via protein-protein interactions with other transcription factors(De Bosscher et al., 2001, 2003; Luecke and Yamamoto, 2005).

**Figure 3:**
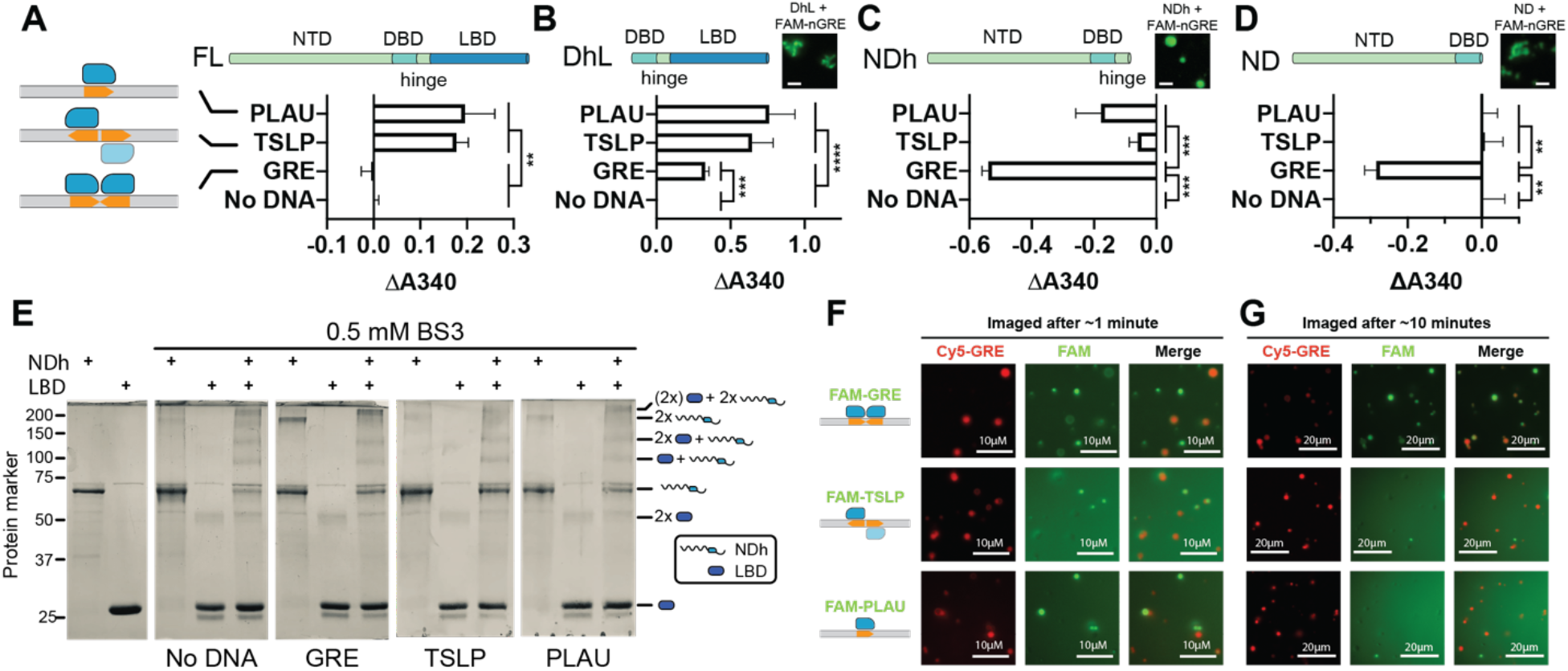
DNA modulates GR phase separation behavior. (**A-D**) Turbidity measurements of different GR constructs with the indicated DNAs. Data is shown as differences in turbidity compared to the “No DNA” condition. Data represents the mean +/− s.e.m. of at least four measurements. One-way ANOVA with Tukey post hoc test for significance of differences between pairs of samples was performed in *Graphpad Prism 8.* Insets: representative images of GR constructs with FAM-labeled DNAs. The length of scale bars is 5 μm. (**E**) Lysine-specific cross-linking confirms that there is an interaction between the LBD and the rest of the protein. (**F**) Representative images of GR condensates with differentially labeled DNAs as indicated. Images were recorded immediately after mixing the samples. (**G**) Representative images of GR condensates with differentially labeled DNAs as indicated. Images were recorded 10 minutes after mixing the samples.

Transactivation via canonical GREs involves recruitment of specific co-activators such as the p160 family of nuclear receptor co-activators (NCoA1, NCoA2, or NCoA3), or MED1(Chen and Roeder, 2007; Chen et al., 2006), a component of Mediator. Conversely, transrepression via repressive DNA elements requires the nuclear receptor co-repressors NCoR1 and NCoR2 as well as histone deacetylases (HDACs)(Surjit et al., 2011). How the selective assembly of co-regulator complexes with GR at specific DNA elements is achieved is not understood.

A growing body of literature is showing that many complex cellular processes occur in biomolecular condensates where functionally related components can be compartmentalized, selectively partitioned, and concentrated(Banani et al., 2016). Condensate formation often involves weak, multivalent interactions between molecules that contain multiple sites for inter- and intramolecular interactions. A prominent feature of many biomolecular condensates is the enrichment of proteins containing large intrinsically disordered regions (IDRs), which provide multivalent, weak intermolecular interactions(Decker et al., 2007; Gilks et al., 2004; Kato et al., 2012; Nott et al., 2015; Reijns et al., 2008). IDRs synergize in condensate formation with additional, specific interactions among folded domains or between folded domains and short peptide motifs(Protter et al., 2018).

Biomolecular condensates have been implicated in most nuclear processes including the regulation of chromosome structure and maintenance, DNA replication and repair, RNA processing, preribosome assembly, and transcription(Sabari et al., 2020). Transcriptional condensates form at specific enhancer foci and contain transcription factors (TFs), transcriptional co-activators, and RNA polymerase II(Boija et al., 2018; Cho et al., 2018; Chong et al., 2018; Fukaya et al., 2016; Hnisz et al., 2017; Sabari et al., 2018; Tsai et al., 2017). A critical characteristic of condensates is their ability to selectively partition proteins of related functions. The determinants that govern selectivity are only beginning to be uncovered. For instance, phosphorylation of its C-terminal IDR regulates the dynamical partitioning of RNA polymerase II between transcriptional and splicing condensates(Guo et al., 2019). Other potential mechanisms that contribute to selective partitioning are the subject of intense ongoing research.

Context-dependent selective partitioning of co-regulators in transcriptional condensates would provide an elegant solution for the assembly of distinct co-regulator complexes with NRs at specific genomic loci. Consistent with this model, GR is located in nuclear foci that have properties of liquid-liquid phase-separated condensates and contain the transcriptional co-activators NCoA2 and MED1, a subunit of the Mediator complex(Stortz et al., 2020; Stortz et al., 2017). The identification of a molecular mechanism underlying the potential selective recruitment of these co-activators into GR condensates, however, requires detailed *in vitro* studies of their phase separation behavior under different functional conditions.

Here, we show that GR forms phase-separated condensates *in vitro,* in a process that requires all parts of the protein – the NTD, DBD, hinge, and LBD. Using GFP-tagged co-regulator IDRs, we show that various known GR co-regulators form droplets *in vitro.* GFP-tagged MED1-, NCoA3-, and G9a-IDRs specifically concentrate in GR condensates, whereas GFP alone or GFP-tagged HDAC2 do not. HDAC2 is known to require other co-regulators for recruitment into GR transcriptional complexes(Bilodeau et al., 2006; You et al., 2013). We then show that GR binding to DNA modulates condensate properties so that co-activator recruitment is enhanced in the presence of activating, but not repressive DNA elements. By introducing mutations into GR and systematic changes in DNA sequences, we determined the molecular mechanism for this behavior using *in vitro* and cell-based assays: proper GR dimerization on canonical GRE DNA, involving a minor groove interaction of a short helix at the DBD C-terminal end, is required for enhanced recruitment *in vitro* and for transcriptional transactivation in cells, but is dispensable for transrepression. These results show that distinct DNA-dependent protein conformations can affect the multivalent interactions in phase-separated condensates controlling the selective recruitment of functionally important interaction partners. Thus, DNA-binding serves as an allosteric selectivity switch governing GR condensate compositional bias. This explains how different DNA elements can cause opposing transcriptional responses in GR-responsive genes. Importantly, the data supports the notion that compositional bias via multivalent, dynamic interactions in nuclear condensates complements known specific, stable interactions between nuclear receptors and co-regulators in the selective assembly of transcriptional complexes.

## Results

### Generation of recombinant full-length GR containing an ancestral LBD

The reconstructed ancestral GR LBD (AncGR2 LBD) has been successfully used to study GR because it reliably recapitulates GR ligand binding, transcriptional responses, and allosteric regulation(Bridgham et al., 2009; Kohn et al., 2012; Liu et al., 2020; Liu et al., 2019; Weikum et al., 2017b). The ancestral GR2 ligand binding domain significantly improves protein expression and stability and shares 79% sequence identity with human GR LBD (Figure S1A). We have generated an intact full-length GR protein containing the ancestral GR2 LBD, which can be purified using a standard bacterial expression system (Figure S1B). The resulting protein is functional in assays for binding to DNA, glucocorticoid ligand, as well as co-regulator peptide (Figure S1C-F).

### GR undergoes reversible liquid-liquid phase separation *in vitro*

Recombinant full-length GR exhibits some unusual behavior. First, solutions above a concentration of approximately 1 mg/mL tend to form strings when extruded out of the thin opening of a pipette tip (Figure 1B). This behavior is reminiscent of pulling fibers from a nylon synthesis reaction and is evidence for extensive intermolecular interactions between the protein molecules. Second, the protein solution becomes turbid when it is removed from ice and placed at room temperature (Figure 1C). This behavior is reversible as the solution turns clear when placed back on ice. When the salt concentration is increased, however, no turbidity is observed at room temperature. Reversible turbidity and sensitivity of this phenomenon to temperature and salt are common characteristics of liquid-liquid de-mixing of proteins in solution. Indeed, fluorescently labelled GR forms well-defined spherical droplets in solution and the number and size of droplets increases with increasing concentration (Figure 1D) confirming that GR forms phase-separated condensates *in vitro*.

GR disordered and ordered domains are required for faithful phase separation behavior IDRs, protein-protein interaction domains, and oligomerization domains are critical drivers of phase separation in many proteins(Elbaum-Garfinkle et al., 2015; Lin et al., 2015; Nott et al., 2015; Pak et al., 2016; Protter et al., 2018; Sabari et al., 2018). To define the contributions of each part of GR to condensate formation, we generated deletion constructs comprising different combinations of its individual domains. Turbidity measurements show that increasing concentrations of GR induce increasing condensate formation (Figure 1E). We find that only full-length protein, but not constructs lacking either the LBD or the N-terminal IDR, undergoes phase separation, showing that intact, full-length protein is required for efficient condensate formation.

Polyethylene glycol (PEG) is a crowding reagent used to more closely simulate the environment in a cell. When placed into a phase separation buffer containing 10% PEG8000 full-length GR forms condensates at approximately 10-fold lower concentrations than in the absence of a crowding reagent (Figure 1F and G). Under these conditions, all GR constructs containing the NTD (ND and NDh) phase separate, whereas a construct lacking the NTD (DhL) does not (Figure 1F). The ND and NDh constructs exhibit progressively reduced phase separation compared to the full-length protein, confirming that the LBD, and also the hinge region, contribute to condensate formation. This is consistent with the weak dimerization of the isolated GR LBD observed previously(Bledsoe et al., 2002). Imaging fluorescently labelled samples of the NDh and ND constructs shows that NDh forms well-defined spherical droplets, whereas ND forms amorphous aggregates (Figure 1G). This observation suggests that this construct exhibits aberrant phase transition behavior akin to the aggregation of disordered proteins in neurodegenerative diseases(Elbaum-Garfinkle, 2019). Thus, the intrinsically disordered hinge region is critical for appropriate intermolecular interactions inducing condensate formation in GR.

Amino acid enrichments in IDRs of proteins forming phase separated droplets tend to point to a mechanism of interaction. The most highly enriched amino acid in the GR N-terminal IDR is serine (Figure S1). Serine residues are required for phase separation of the IDR of MED1(Sabari et al., 2018). To test if GR phase separation requires serines we mutated all 61 serines within the N-terminal IDR to alanines. The resulting mutant behaves very similar to the corresponding wild type protein showing that GR phase separation is not driven by serine residues (Figure 1F).

Next, we tested if deletion constructs are miscible with condensates of full-length GR (Figure 1H). Imaging of differentially labelled proteins (Cy5-GR and Cy3-labelled deletion constructs) shows that the DhL construct, which does not contain the N-terminal IDR, does not get recruited into GR droplets. In contrast, the NDh construct, at a concentration at which it does not form droplets alone, is enriched on the surface of GR droplets, but does not efficiently diffuse into the droplets’ interior. Thus, this construct can undergo phase separation alone and gets concentrated on the surface of GR droplets, but without an LBD will not efficiently mix with droplets formed by full-length GR.

Together these results show that GR undergoes phase separation reversibly and that all of its domains, structured and non-structured, cooperate and contribute to the faithful implementation of this process, i.e. efficient condensate formation, prevention of aggregation, and miscibility of droplets. This highlights the fact that structured domains play important roles in regulating the phase transition of proteins.

### Known GR transcriptional co-regulators exhibit phase separation characteristics *in vitro*

GR activates and represses gene expression by recruitment of various co-regulator proteins and many of these proteins are enriched in IDRs. The MED1 subunit of Mediator uses its long IDR to phase separate *in vitro(Boija et al., 2018; Sabari et al., 2018; Shrinivas et al., 2019)* (Figure 2A). To test if other known GR co-regulators exhibit phase separation behavior *in vitro,* we expressed GFP-fusions of the IDRs of the co-activator NCoA3, the co-repressor HDAC2, and the histone methyltransferase G9a, which is primarily a co-repressor but also displays co-activating activities towards glucocorticoid-responsive genes (Figure 2A)(Bittencourt et al., 2012). When placed into phase separation buffer containing PEG8000, all proteins formed well-defined spherical droplets at micromolar concentrations (Figure 2B). Notably, HDAC2 only showed evidence for droplets at concentrations above 10 μM whereas the other proteins formed droplets at 1 μM. These results confirm that the IDRs of known GR co-regulators can undergo phase separation *in vitro.*

**Figure 2:**
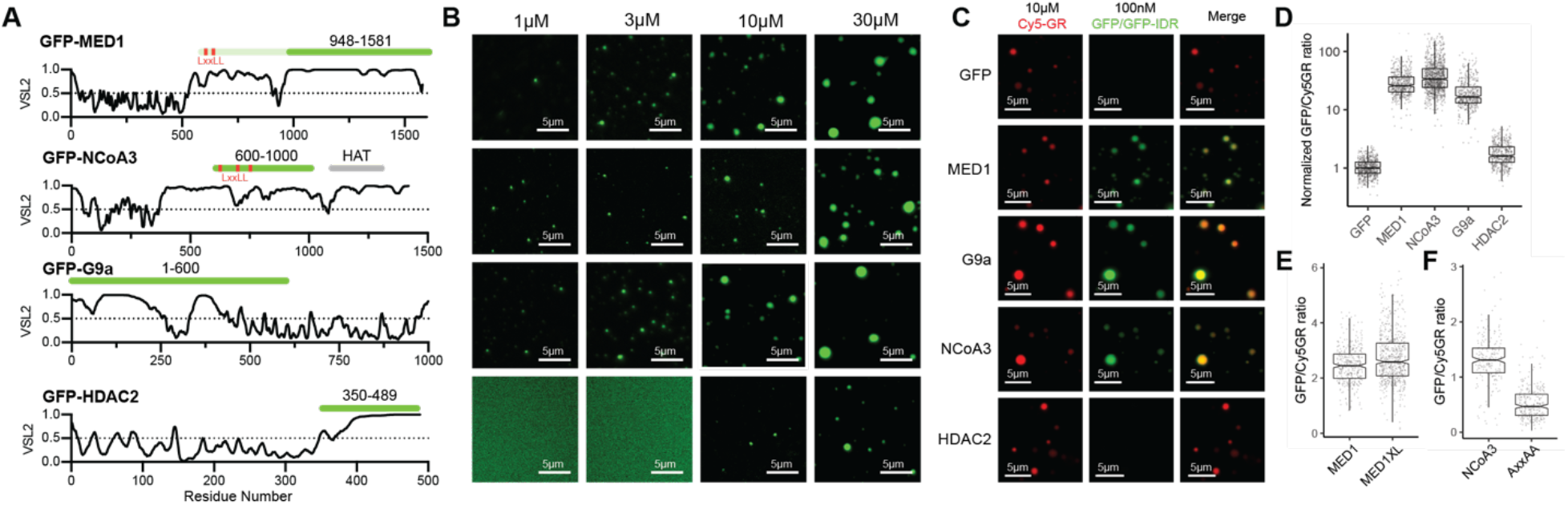
Co-regulator IDRs undergo phase separation and specifically concentrate in GR condensates. **(A)** VSL2 disorder scores for known GR co-regulators. Green bars represent GFP-tagged IDRs used in this study. The bar in a lighter green shade in MED1 represents the additional sequence included in a second, longer construct, which covers its two LxxLL motifs (MED1XL). The location of the HAT domain of NCoA3 is highlighted with a grey bar. **(B)** All co-regulator IDRs form phase separated droplets at increasing concentrations. HDAC2, contains the shortest IDR and induces droplets only above 10μM. **(C)** GR condensates specifically concentrate the IDRs of MED1, G9a, and NCoA3, but not GFP alone or the IDR of HDAC2. **(D)** Quantification of fluorescence signals in condensates shown in (C). Every dot represents a detected droplet. **(E)** Quantification of the recruitment of MED1 constructs with and without LxxLL motifs into GR condensates. **(F)** Quantification of the recruitment to GR condensates of wild-type NCoA3 and a variant in which LxxLL motifs are mutated to AxxAA.

### Known GR transcriptional co-regulators concentrate in GR condensates

One of the functional properties of membrane-less organelles is the enrichment and exclusion of specific molecules in order to increase local concentrations of functionally related proteins or to prevent unwanted interactions with unrelated proteins. Enrichment of several transcription factors has been shown for condensates of MED1 including the estrogen receptor, a member of the nuclear hormone receptor family(Boija et al., 2018). To test if GR condensates can specifically enrich MED1 we imaged GR droplets in the presence of GFP-MED1 at concentrations at which it does not phase separate alone showing that that GFP-MED1-IDR is efficiently recruited into GR droplets while GFP alone is not (Figure 2C).

To test if the IDRs of other known co-regulators exhibit the same behavior, we repeated this experiment with the GFP-tagged IDRs described above (Figure 2 C and D). The IDRs of NCoA3 and G9a are specifically concentrated in GR condensates, whereas the IDR of HDAC2 is not. To account for the reduced efficiency of droplet formation of the HDAC2 IDR we increased the GFP-HDAC2-IDR concentration to 1 μM. Even at the increased concentration GFP-HDAC2-IDR is not substantially enriched in GR condensates compared to GFP alone (Figure S2A). This is consistent with the observation that HDAC2 interaction with GR requires other co-regulators such as the Brg1 ATPase subunit of the Swi/Snf complex(Bilodeau et al., 2006) or nuclear receptor co-repressor NCoR1(You et al., 2013). Together, these results show that GR condensates specifically enrich or exclude known transcriptional co-regulators via their IDRs *in vitro,* and its behavior is consistent with known interactions established in the literature.

The NCoA3-IDR used here (residues 600-1000) contains three LxxLL motifs, while the entire G9a protein does not contain any (Figure 2A). There are two LxxLL motifs present in MED1, however, the MED1-IDR construct used here does not cover these (Figure 2A). This shows that a stable interaction between GR and co-regulators is not required for efficient enrichment in GR condensates. To further test the role of the LxxLL motif we measured recruitment of two additional constructs. We mutated the motifs in NCoA3 to AxxAA and generated a longer MED1 construct covering its LxxLL motifs, MED1XL(Sabari et al., 2018)(Figure S2). GFP-MED1XL was efficiently recruited to GR droplets to a similar degree as GFP-MED1-IDR (Figure 2F). The NCoA3 AxxAA mutant was also enriched in GR condensates, but to a significantly lower degree than the wild-type IDR (Figure 2G). These results show that recruitment of co-regulators into GR condensates is driven by a combination of multivalent, dynamic interactions via IDRs as well as stable, specific interactions via LxxLL motifs and the AF2. Recruitment of MED1 appears be driven strongly by IDR interactions, whereas NCoA3 recruitment is significantly enhanced by the presence of intact LxxLL motifs.

### DNA binding modulates GR condensate properties

The transcriptional activity of GC-responsive genes is determined in part by the type of GR response element found in their associated regulatory DNA regions. The DNA response element impacts which co-regulators – co-activators or co-repressors – are recruited into agonist-bound GR transcriptional complexes. Structural studies have determined the different arrangements of GR on these DNA response elements(Weikum et al., 2017a). How these different arrangements induce opposite transcriptional responses, however, is not understood. We asked if DNA binding induces changes in GR phase separation behavior that might provide some clues about selective co-regulator interactions.

We characterized phase separation behavior of GR bound 40 bp DNA duplexes representing different types of GR response elements: a canonical GRE, an inverted repeat GRE found in the TSLP promoter(Hudson et al., 2013; Surjit et al., 2011), or the cryptic half-site found in the κBRE of the PLAU promoter. The degree of condensate formation was determined by measuring the solutions’ turbidity. We observed small, but significant increases in turbidity when TSLP or PLAU DNA were added to full-length GR, but not upon addition of GRE DNA (Figure 3A). Dimerization commonly is a potent driver of phase separation, so it is surprising that a canonical GRE, which causes co-operative GR binding and dimerization, did not result in turbidity changes whereas a half-site containing DNA (PLAU) did. What is more, in GR constructs that do not contain an LBD, phase separation is greatly reduced upon dimerization on GRE-containing DNA (ND and NDh; Figure 3C and D). Binding of GR to TSLP or PLAU DNA increases turbidity compared to GRE DNA for all constructs tested.

The DhL construct is highly sensitive to DNA binding and interaction with DNA causes the protein to aggregate rather than form spherical droplets characteristic of phase separated condensates (Figure 3B). This aberrant phase transition behavior is a sign of enhanced intermolecular interactions and may provide useful mechanistic insight into GR condensate formation. For instance, it suggests that in the full-length protein the N-terminal IDR protects against unfavorable interactions present in the DhL construct. Unfavorable interactions are most likely mediated by the LBD since canonical GRE binding to constructs missing the LBD (NDh or ND) reduces, rather than increases, turbidity compared to the DNA-free condition (Figure 3C and D). These observations suggest a direct interaction between the NTD and LBD in the full-length protein. Indeed, the addition of a chemical cross-linker to a solution of the NDh construct and the LBD identifies an interaction between the two (Figure 3E).

Together these data show that DNA binding induces allosteric changes in GR that modulate condensate formation. In the full-length receptor, interactions between the NTD and LBD are balancing GR self-association to ensure functional condensate formation. These observations underscore the above-mentioned complexities in the interplay between the different structured and unstructured domains of GR. The fact that activating and repressive GR DNA-binding elements have opposing effects suggests that the known, distinct GR binding modes on these DNAs are communicated via the NTD and LBD and ultimately cause differences in condensate properties.

### DNA-binding induces selective partitioning in GR condensates

The substantial DNA-induced effects on GR phase transitions suggest profound underlying differences in the interactions driving GR condensate formation. One of the features of condensates thought to drive compositional bias of components is the particular milieu formed by amino acid side chains intermingling dynamically via conventional intermolecular interactions(Sabari et al., 2020). This led us to ask if DNA binding causes changes in the molecular interaction milieu of GR condensates that could lead to selective partitioning.

To investigate this question, we tested if condensates of GR-DNA complexes are readily miscible with each other. First, GR was bound to Cy5-labelled GRE DNA. In parallel, GR was bound to different fluorescein amidite(FAM)-labelled DNA elements (GRE, TSLP, and PLAU). Condensate formation was then induced before mixing Cy5-GRE samples with the individual FAM-DNA samples (Figure 3F-G). It is important to note here that we were imaging fluorescently labelled DNAs and not GR itself. GR interactions with DNA are thought to be highly dynamic(Fletcher et al., 2002; Nagaich et al., 2004; Presman et al., 2016) so it is likely that DNA is repeatedly unbinding from and re-binding to GR during the experiment. Imaging showed that GR-GRE condensates readily exchanged with each other and after 10 minutes Cy5-GRE and FAM-GRE signals were randomly distributed between GR droplets (Figure 3G and Supplementary Figure 3). Droplets containing FAM-TSLP and FAM-PLAU, however, were only visible when imaged immediately after mixing the samples and the FAM-signal was mostly diffuse after a 10-minute incubation, with only very few FAM-containing condensates left (Figure 3F and G). In fact, FAM-TSLP and FAM-PLAU were excluded from Cy5-GRE containing droplets as evidenced by shadows in the FAM-signal (Figure 3G).

Images taken immediately after mixing the samples showed that Cy5-GRE was initially accumulating on the surface of GR condensates containing FAM-TSLP or FAM-PLAU, but not vice versa (Figure 3B). Conversely, Cy5-GRE did not associate with the surface of FAM-GRE containing droplets. FAM-labeled DNAs were added in slight excess in order to saturate GR in those samples before mixing. As a result, FAM-GRE was concentrating on the surface of Cy5-GRE containing droplets immediately after mixing. This was not observed for FAM-TSLP or FAM-PLAU. This shows that repressive DNAs are excluded from GR condensates formed by GR bound to an activating DNA, whereas GR complexed with differentially labeled activating DNAs (FAM/Cy5-GREs) dynamically distribute between the solution and condensates.

We conclude that DNA-binding changes the milieu of potential intermolecular interactions present in GR condensates. This causes immiscibility of GR condensates containing activating versus repressive DNAs and suggests that GR is capable of forming physically separate activating and suppressive condensates *in vivo.*

### GRE DNA binding increases MED1-IDR recruitment to GR condensates

Next, we assessed if changes induced by DNA also affect co-regulator recruitment. We assembled GR-DNA complexes, formed condensates in the presence of 100nM GFP-MED1-IDR, and quantified these by fluorescence microscopy (Figure 4A and B). The results revealed two main changes in GR condensates upon DNA binding. First, binding to GRE DNA increases the relative amount of GFP-MED1-IDR accumulating in GR condensates compared to TSLP or PLAU DNA or in the absence of DNA (Figure 4B). Second, GFP-MED1-IDR formed puncta on the surface of GR condensates in the presence of TSLP or PLAU DNA suggesting that it phase-separated from within GR condensates under these conditions (Figure 4A). Consistent with this notion, this behavior was more pronounced when the MED1-IDR concentration was increased to 3 μM (Figure 4C). At this concentration puncta appeared greater in size and coalesced into larger, continuous structures around the GR condensates. At 100 nM, MED1-IDR only induced very few and small puncta in GR condensates containing GRE DNA. However, clear puncta appeared at 3 μM GFP-MED1-IDR which is consistent with phase separation within GR condensates once a critical concentration is reached.

**Figure 4:**
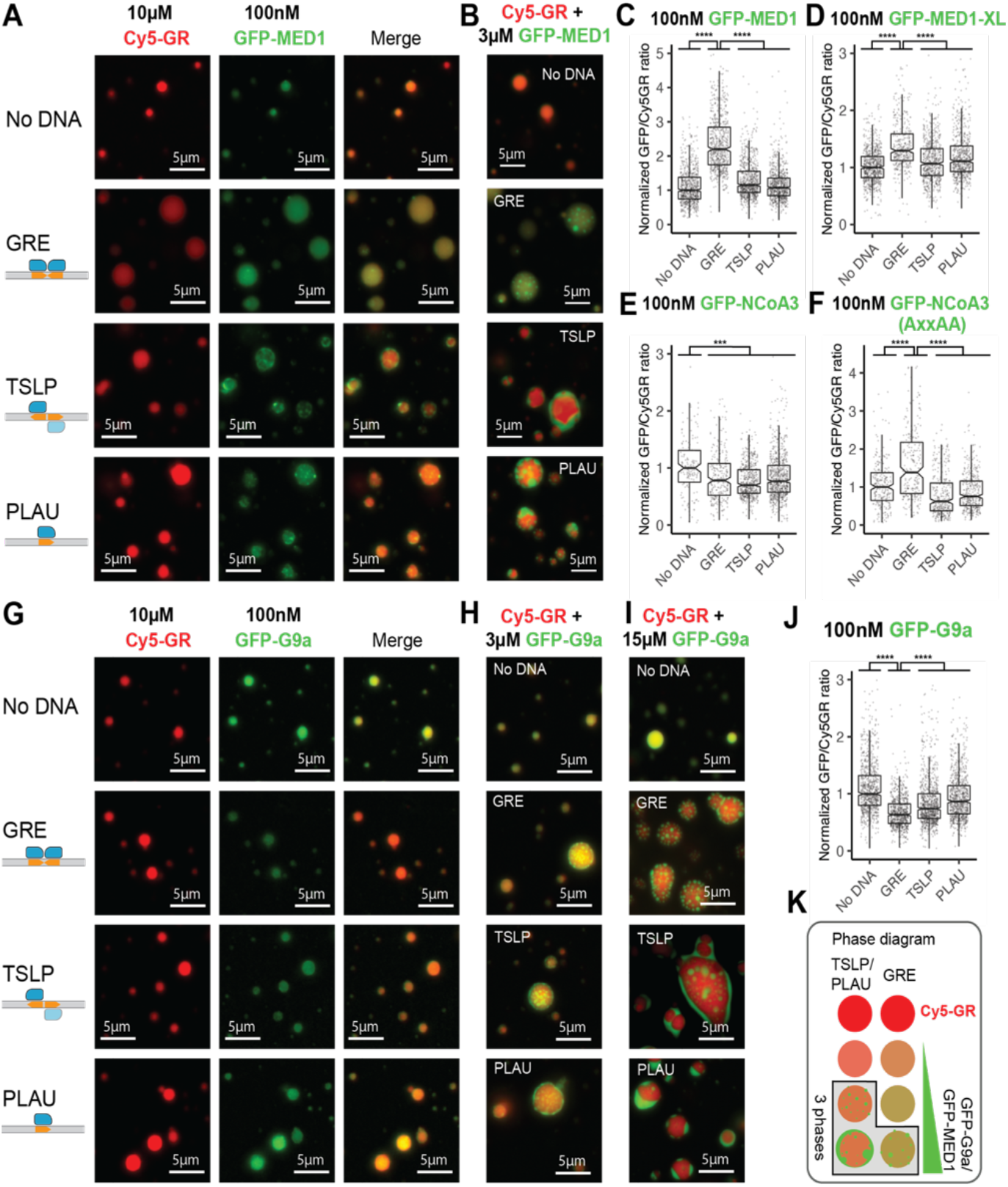
DNA binding modulates co-regulator recruitment consistent with their activating and repressive functions. (**A**) Representative images of Cy5-labeled GR condensates with GFP-MED1 at low concentration (100 nM) and the indicated DNAs. (**B**) Representative images of Cy5-labeled GR condensates with GFP-MED1 at high concentration (3 μM) and the indicated DNAs. (**C**) Quantification of fluorescence intensity ratios in the GFP- and Cy5-channels for condensates shown in (A). Each dot represents one detected droplet. Data was normalized to the “No DNA” condition. (**D-F**) Quantification of fluorescence intensity ratios in the GFP- and Cy5-channels for Cy5-labelled GR condensates with the indicated GFP-tagged proteins. Each dot represents one detected droplet (Representative images are shown in Supplementary Figure 4). Data was normalized to the “No DNA” condition. (**G-I**) Representative images of Cy5-labeled GR condensates with GFP-G9a at the indicated concentrations. (**J**) Quantification of fluorescence intensity ratios in the GFP- and Cy5-channels for Cy5-labelled GR condensates with GFP-G9a at 100nM (images shown in (G)) Each dot represents one detected droplet. (**K**) Phase diagram describinbg the transition from two-phase to three-phase behavior of Cy5-GR condensates in the presence of DNA and co-regulators. (C-F and J) At least 5 images were analyzed for each sample. P values were calculated using Welch’s t-test. ****: adjusted p value < 0.0001.

These results show that DNA-binding modulates the miscibility of GR condensates with MED1-IDR. The co-activator MED is selectively recruited in the presence of GRE DNA, consistent with transcriptional activation mediated by this sequence. Changes in the recruitment of MED1-IDR appears to involve changes in the miscibility of this co-regulator within GR condensates resulting in three-phase behavior when miscibility is reduced in the presence of negative response elements such as inverted repeats (TSLP) or GR half-sites (PLAU) (Figure 4K).

### DNA-dependent recruitment of co-activators is modulated by LxxLL motifs

Next, we investigated the role of LxxLL motifs in the selective enrichment of co-activators in GR condensates. GFP-MED1-XL, which contains two LxxLL motifs, was selectively enriched in GRE-containing GR condensates, albeit to a somewhat lesser extent than the shorter MED1 construct (Figure 4D). GFP-MED1-XL did not exhibit any three-phase behavior (Figure S4C). The recruitment of GFP-NCoA3, which contains three LxxLL motifs, was slightly decreased by the addition of all DNAs with no evidence for three-phase behavior (Figure 4E; Figure S4A and B). The AxxAA mutant of NCoA3, however, exhibited enhanced recruitment in the presence of GRE DNA (Figure 4F). These observations are consistent with the trends observed for the role of LxxLL motifs in the absence of DNA (Figure 3E and F). It appears that multivalent, dynamic interactions mediated by the IDRs of NCoA3 and MED1 are increased when GR is bound to GRE DNA. In NCoA3 stable interactions via the LxxLL motifs are more important for recruitment (Figure 2F) so that they overcome the effect of enhanced interactions via IDRs, at least in the *in vitro* setting used here. It is possible that the multivalent interactions mediated by the NCoA3 IDR play a more prominent role in condensate recruitment in the cellular environment, where there is competition for co-regulator binding(Kamei et al., 1996; McKenna and O’Malley, 2002).

### G9a recruitment to GR condensates is reduced by DNA binding

The co-repressor G9a does not contain any LxxLL or CoRNR box motifs and recruitment of GFP-G9a at a concentration below its own phase separation threshold is reduced when GR is bound to DNA (Figure 4G). Condensates of GR complexed with GRE DNA recruit GFP-G9a less efficiently than condensates of GR bound to repressive DNAs (Figure 4J). At higher concentrations of GFP-G9a (3 μM) we observed clear three-phase behavior with GFP-G9a droplets on the surface of GR condensates in the presence of DNA, but not with GR alone (Figure 4H). At even higher concentrations (15 μM) GFP-G9a droplets grew larger and coalesced around the GR condensates (Figure 4I) similar to MED1. These results confirm that DNA binding modulates recruitment of co-regulators consistent with their activating and repressive functions.

Multi-phase separation is a common phenomenon in biological systems with many components(Jacobs and Frenkel, 2017; Riback et al., 2020; Shin and Brangwynne, 2017). Examples include paraspeckles(West et al., 2016), PML bodies(Lang et al., 2010), stress granules(Jain et al., 2016), the nucleolus(Feric et al., 2016), and Cajal bodies(Gall et al., 1999). The separation of multiple non-coalescent phases is considered a mechanism for tuning condensate composition and to facilitate sequential reactions in multi-step processes such as ribosome assembly(Feric et al., 2016; Riback et al., 2020). It appears that GR may utilize this mechanism to control the composition of transcriptional condensates by excluding distinct co-regulators depending on its DNA-binding mode (Figure 4K).

### Helix H3 binding on GRE DNA is required for enhanced recruitment of MED1-IDR to GR condensates and transcriptional transactivation

GR binding on DNA via the DBD involves specific interactions through conserved residues in the DNA-reading helix and additional interactions contributed by a short helix at the DBD C-terminus. This helix 3 (H3) is highly positively charged, is flexible in the absence of DNA(Frank et al., 2018), and lines up along the DNA minor groove 3 bp on the distal side of each half-site in a stable conformation when bound to GRE DNA(Meijsing et al., 2009). Because it connects the DBD to the short hinge and the LBD, we have previously hypothesized that the orientation of H3 along the minor groove may serve to constrain the LBD in an active conformation competent for dimerization on a GRE, but not on other DNA elements(Frank et al., 2018). To test if H3 DNA-binding and the potential resulting LBD dimerization explain the observed differences in GR phase transitions and co-regulator recruitment we designed DNAs of equal length, in which the GRE is moved along the length of the duplex in a stepwise manner (Figure 5A). As the GRE approaches the edge, the distal H3 will not be stabilized in the DNA minor groove and the proposed LBD dimerization will be disfavored.

**Figure 5:**
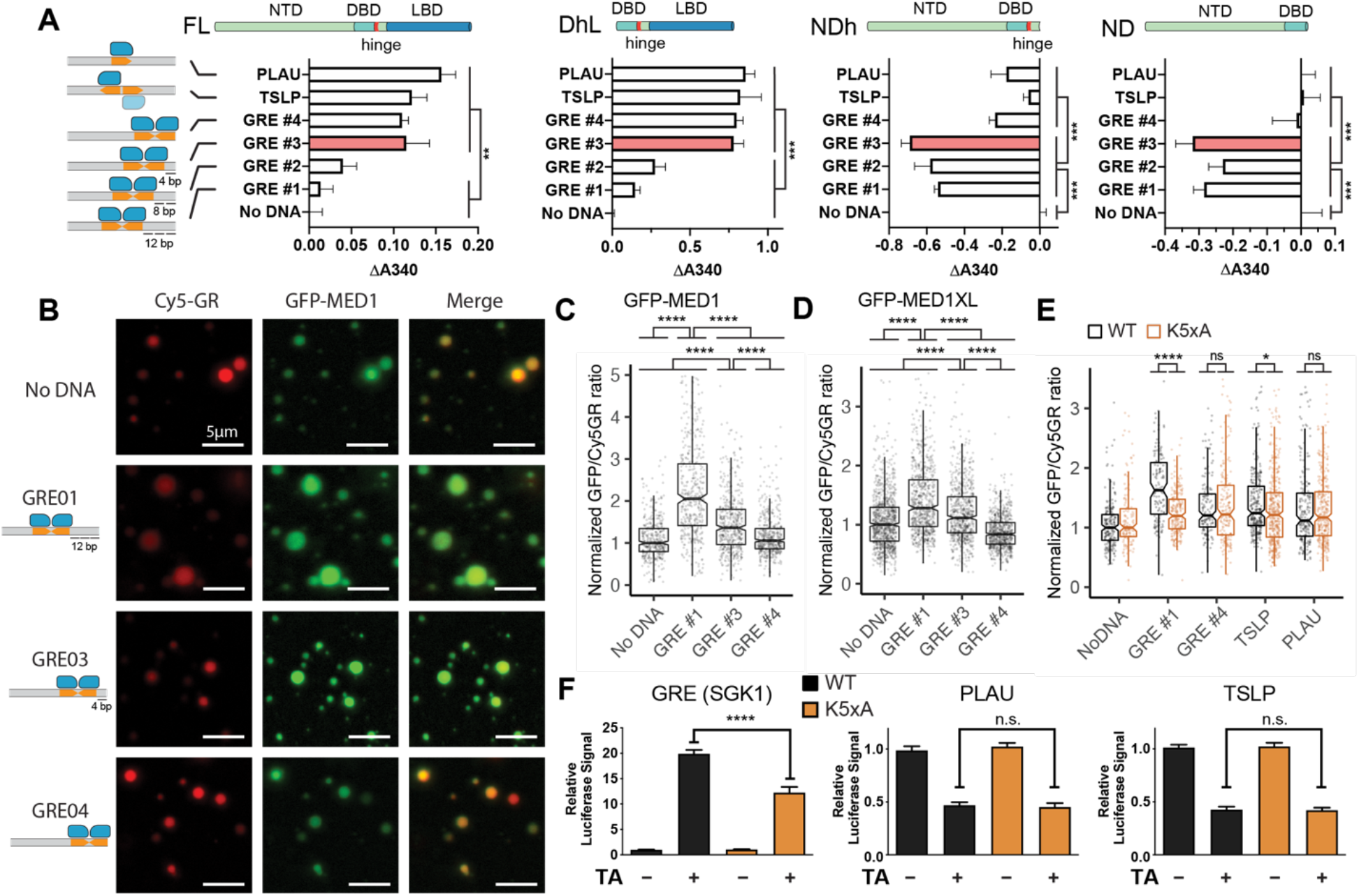
Helix H3 binding on GRE DNA is required for enhanced co-activator recruitment and transcriptional transactivation. (**A**) Turbidity measurements of different GR constructs with the indicated DNAs. Data is shown as differences in trubidity compared to the “No DNA” condition. Data represents the mean +/− s.e.m. of at least four measurements. One-way ANOVA with Tukey post hoc test for significance of differences between pairs of samples was performed in *Graphpad Prism 8.* (**B**) Representative images of Cy5-labeled GR condensates with GFP-MED1 at low concentration (100 nM) and the indicated DNAs. (**C**) Quantification of fluorescence intensity ratios in the GFP- and Cy5-channels for condensates shown in (B). Each dot represents one detected droplet. Data was normalized to the “No DNA” condition. (**D**) Quantification of fluorescence intensity ratios in the GFP- and Cy5-channels for Cy5-labelled GR condensates with GFP-MED1XL at 100nM. Representative images are shown in Supplementary Figure 5. Each dot represents one detected droplet. Data was normalized to the “No DNA” condition. (**E**) Quantification of fluorescence intensity ratios in the GFP- and Cy5-channels for condensates of Cy5-labelled wild type GR or the K5xA mutant with GFP-tagged GFP-MED1. Representative images are shown in Supplementary Figure 5. Each dot represents one detected droplet. Data was normalized to the “No DNA” condition. (**F**) Reporter gene assays showing that the GR K5xA mutant is impaired in its ability to trans-activate (SGK1 GRE), but not trans-repress (TSLP and PLAU). One-way ANOVA with Tukey post hoc test for significance of differences between pairs of samples was performed in *Prism.* (C-E) At least 5 images were analyzed for each sample. P values were calculated using Welch’s t-test.

GRE #3, in which the GRE site is 4 bp from the edge, increases phase separation in full-length GR and DhL to similar extent as TSLP or PLAU (Figure 5A). In ND or NDh, which do not contain the LBD, however, this DNA behaves the same as GRE #1. DNA end-fraying in GRE #3 likely prevents H3 stabilization, which explains the effects on full-length GR and DhL. We conclude that H3 binding to DNA is critical for changes in the phase transition behavior of GR when bound to GRE as opposed to TSLP or PLAU DNA and that the LBD is required for this behavior. GRE #4, which contains the GRE site at the edge of the duplex with no additional flanking base pairs, induces increased phase separation similar to TSLP and PLAU in all GR constructs, including NDh and ND. The most likely explanation for this behavior is that end-fraying abolishes the distal half-site of the GRE so that this DNA behaves like PLAU, i.e. a single half-site.

Helix 3 binding is also required for the observed enhanced recruitment of MED1-IDR to GR condensates in the presence of GRE DNA (Figure 5B and C). GRE #3 and GRE #4 cause gradually less incorporation of GFP-MED1-IDR into GR condensates compared to GRE #1. MED1XL exhibited the same behavior (Figure 5D). Next, we mutated the lysine residues within helix H3 (“K5xA”) to prevent H3 binding in the minor groove. As expected, GR(K5xA) exhibits reduced overall affinity for DNA (Figure S5). However, this mutant retains increased affinity towards GRE DNA compared to PLAU and TSLP consistent with the ability to cooperatively dimerize on GRE DNA via interactions through the dimerization loop. Quantitative imaging showed that GR(K5xA) does not exhibit enhanced recruitment of MED1-IDR to condensates in the presence of GRE DNA (Figure 5E) confirming that H3 binding is critical for selective co-regulator recruitment.

Since dimerization only occurs on activating, but not repressive DNA elements, we hypothesized that mutation of H3 would affect transactivation, but not transrepression of GR-responsive genes. To test the transcriptional activity of GR(K5xA) in cells we performed luciferase reporter assays. A reporter driving transactivation via canonical GRE motifs (SGK1) showed reduced transactivation activity of GR(K5xA) compared to wild type GR (Figure 5F). Transrepression via TSLP or PLAU elements, however, was not affected by the lysine mutations confirming that proper dimerization via H3 binding is required for transactivation, but not transrepression.

These results show that DNA-dependent changes in GR phase transitions correlate with the selective recruitment of GFP-MED1-IDR to GR condensates and require proper GR dimerization including the minor groove interaction via H3. This provides a molecular mechanism for the DNA sequence-dependent transcriptional transactivation activity of GR mediated by canonical GREs.

## Discussion

A growing body of literature is showing that phase separation is a driving force in the transcriptional activation of genes(Boija et al., 2018; Cho et al., 2018; Chong et al., 2018; Guo et al., 2019; Sabari et al., 2018; Sabari et al., 2020; Shrinivas et al., 2019). The proteins involved in this process – transcription factors, transcriptional coactivators, and RNA polymerase – use their IDRs to form phase-separated condensates that enhance transcription. Many transcription factors are bifunctional, containing a folded DBD for genomic localization and a disordered activation domain(Sigler, 1988) that weakly associates with co-regulators in dynamic nuclear condensates(Boija et al., 2018). NRs are multifunctional with an additional ligand-regulated domain and a disordered hinge connecting the DBD to the LBD. We show here how GR, a prominent member of the NR family, utilizes this additional layer of regulatory potential to up- or down-regulate the transcription of target genes in response to distinct DNA stimuli.

We show that GR forms phase-separated condensates *in vitro* and that each individual domain contributes to this behavior. While stable, folded domains have been shown to contribute to phase separation in other proteins, the complex cooperation of multiple domains in GR is unique. The interplay of ordered and disordered regions is further highlighted by the differential responses of different GR constructs to the binding of activating and repressive DNA elements (Figure 3A and 5A). These interactions modulate condensate properties in two important ways. Droplets of GR dimerized on GRE DNA are not miscible with GR bound to repressive DNA sequences so that physically separate activating and repressive droplets can be formed (Figure 6A). Activating droplets containing GRE DNA selectively recruit the IDRs of co-activators (MED1 and NCoA3) and exclude co-repressor IDRs (G9a). This provides a convenient mechanism for the DNA sequence-dependent control of transcriptional up- and down-regulation of genes by GR at specific genomic loci.

**Figure 6:**
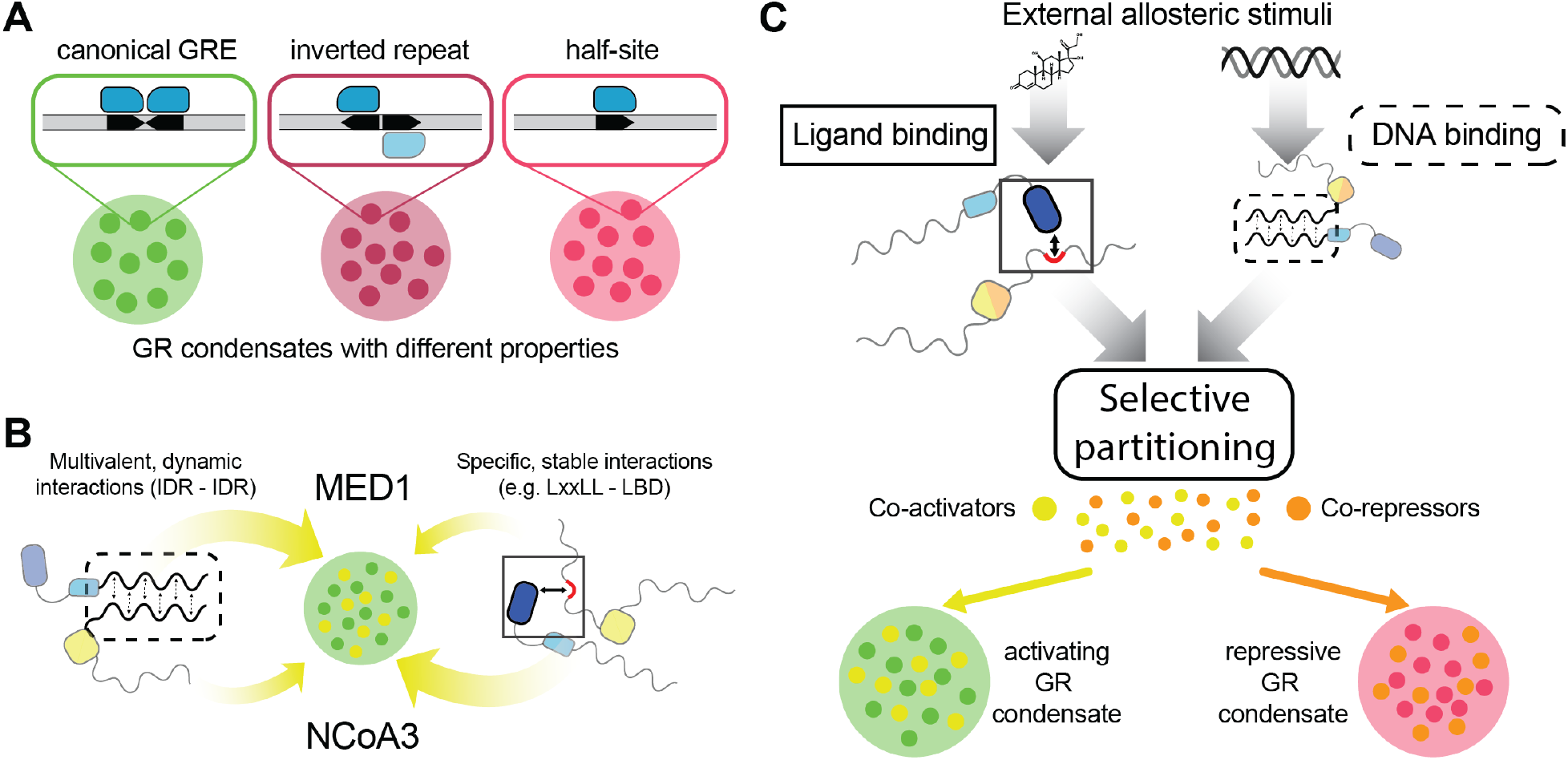
Model for the selective partitioning of co-regulators in GR condensates. (**A**) Distinct DNA binding elements change the properties of GR condensates. (**B**) IDR-mediated, multivalent interactions and stable, specific interactions between LxxLL motifs and the LBD contribute to varying degrees to the recruitment of co-regulators to GR condensates. Multivalent interactions dominate in MED1, whereas stable interactions dominate the recruitment of NCoA3. This explains the sensitivity of MED1 to DNA-induced changes of GR condensate properties and NCoA3’s strong dependence on LxxLL motifs. (**C**) A model for the selective partitioning of co-regulators among GR condensates. DNA and ligands are external stimuli that affect selective partitioning of coregulators independently. Ligand-binding allosterically modulates specific interactions via LxxLL motifs and the LBD, whereas DNA binding allosterically modulates multivalent interactions via IDRs.

We identify a molecular mechanism for how selective partitioning of co-regulators is controlled by DNA binding. This phenomenon requires dimerization of GR on DNA and a minor-groove interaction of helix H3. Preventing H3 association with DNA by either moving the GR binding site to the edge of the DNA or by mutation of positively charged residues in H3 abolishes selective recruitment of co-regulators. Cellular assays show that mutations of H3 prevent GR-mediated transactivation, but not transrepression, consistent with GR dimerization on activating, but not repressive DNA elements. Interestingly, lysine residues in H3 are acetylated by the histone acetyltransferases CLOCK and BMAL1, which inhibits GR transactivation(Nader et al., 2009). Our results suggest that this inhibition may result from consequences of H3 binding on the DNA. Another study showed that deacetylation of K494 and K495 by HDAC2 is required for GR-dependent repression of NF-κB signaling by controlling the interaction between GR and NF-κB (Ito et al., 2006). Together, these observations and our data show H3 lysine residues play a critical role in regulating GR signaling activities by affecting co-regulator recruitment.

How does DNA binding cause changes in condensate selectivity? One of the features thought to control selective partitioning between condensates is the milieu of amino acid side chains available for dynamic, multivalent interactions (Sabari et al., 2020). Our results suggest that DNA binding coordinates interactions between the disordered NTD and the folded LBD. This may remove the sequence surrounding the site of interaction from the pool of amino acids available for dynamic interactions that are responsible for condensate formation and compositional bias. Another source for compositional bias is specific, stable interactions. We show that the contributions of LxxLL motifs to the selective partitioning in GR condensates differs between co-regulators. MED1 is efficiently enriched even in the absence of any LxxLL motifs and the addition of two LxxLL motifs does not increase MED1 concentration in GR droplets showing that MED1 recruitment is dominated by multivalent, dynamic interactions mediated via its IDR (Figure 6B). In contrast, NCoA3 recruitment is controlled by specific, stable interactions since its concentration in GR droplets is highly dependent on LxxLL motifs.

From these data a model emerges in which specific, stable interactions via LxxLL motifs and multivalent, dynamic interactions represent two independent mechanisms for co-regulator recruitment by GR, and likely by NRs in general (Figure 6C)(Protter et al., 2018; Sabari et al., 2020). Each mechanism may contribute to a different degree in the recruitment of a particular NR co-regulator to a nuclear condensate as observed for MED1 and NCOA3 here (Figure 6B). Additionally, each mechanism can selectively and independently be controlled by external stimuli. NR-binding peptide motifs – NR boxes and CoRNR boxes – have varying affinities for NRs at the AF2 surface. Ligand binding in the LBD modulates the conformation of the AF2, which controls the relative affinities of these motifs. As a result, different ligands induce differential co-regulator interaction profiles in NRs(Desmet et al., 2017). Analogously, we show here that co-regulator IDRs have varying propensities to concentrate in GR condensates via multivalent interactions with the GR N-terminal IDR. DNA binding modulates the condensate interaction milieu, most likely by changing the amino acids available for interaction with co-regulator IDRs. Importantly, different DNA elements induce differential effects on condensates, which controls the recruitment of co-activators and co-repressors. In analogy to the allosteric control of AF2 via ligand binding, we show that DNA acts as an allosteric selectivity switch by controlling the AF1. This model provides a possible explanation for the tissue- and gene-specific assembly of distinct transcriptional complexes. Many co-regulators appear to function as components of large, multi-protein complexes, yet the number of potential coregulators exceeds the capacity for direct interaction by a single receptor (Glass and Rosenfeld, 2000). We propose that context-dependent assembly of NR complexes is a consequence of compositional bias in nuclear receptor condensates, which can be modulated by a number of factors: (i) The intrinsic propensity of co-regulator IDRs for selective partitioning into a particular condensate, which may be DNA-dependent for some NRs, (ii) ligand-dependent co-regulator interaction profiles, (iii) tissue-specific expression of co-regulators, and (iv) the presence of other transcription factors in the local environment around a receptor binding site. The environment here accounts for the presence of other transcription factors that bind nearby genomic regions. Depending on the properties of their IDR these TFs may be present in NR condensates and thus potentially change condensate interaction milieu to impact co-regulator recruitment. Together these factors may provide the level of selectivity required for the observed tissue- and gene-dependent assembly of nuclear receptor transcriptional complexes.

## Acknowledgements

This work was supported by a W.M. Keck Foundation Medical Research Grant. F.F. was supported by a *Winship Invest$* pilot grant. E.A.O was supported by grant R01DK115213 from the NIH National Institute of Diabetes and Digestive and Kidney Diseases.

## Methods and Materials

### Plasmids

Protein expression plasmids are based on the previously published pJ411 bacterial expression vector containing the NTD and DBD (residues 1-525), which is named NDh here (Li et al., 2017). Full-length GR containing the ancestral GR2 LBD was cloned by inserting the GR2 LBD into this vector. Briefly, the pJ411 plasmid was PCR amplified to generate a linear DNA fragment containing the plasmid backbone and the NDh construct without a STOP codon. The GR2 LBD was amplified by PCR using oligonucleotide primers to insert overlaps with the amplified plasmid DNA. The two fragments were combined using Gibson assembly. The ND construct was generated by PCR-amplification of the pJ411 plasmid backbone (without any GR sequence) and insertion of a fragment containing GR residues 1-487.

The DhL construct was cloned into the pSmt3 plasmid(Mossessova and Lima, 2000) using Gibson assembly to generate a His6-SUMO-tagged fusion protein.

The bacterial expression construct for GFP-MED1 in the pETEC-GFP plasmid(Sabari et al., 2018) was generously provided by Richard A. Young’s lab. GFP-MED1XL was generated by first amplifying the MED1XL sequence from pETEC-mCherry-MED1XL (provided by Richard A. Young’s lab)(Sabari et al., 2020) and inserting it into the pETEC-GFP plasmid linearized with BamHI and EcoRI restriction enzymes using a Gibson assembly. NCoA3 (residues 600-1000), G9a (residues 1-600), and HDAC2 (residues 350-489) were inserted into pETEC-GFP using Gibson assembly.

The GR K5xA mutant (492-KTKKKIK-498 to 492-ATAAAIA-498) was generated by first PCR amplifying two fragments of GR (residues 1-491 and 499-777) using primers to insert the mutated residues 492 to 498 so that the two fragments overlapped in the mutated region. These fragments were then Gibson assembled into pJ411.

The LxxLL to AxxAA mutant of NCoA3 was generated using the same approach. Three LxxLL motifs and an additional LxxIL motif were mutated to AxxAA (Supplementary Figure 2). Five PCR fragments were amplified with overlaps containing the four mutated AxxAA sequences and Gibson assembled in a single step into the pETEC-GFP plasmid.

### Protein expression and purification

Proteins were expressed in *E. coli* BL21 DE3. Cells were grown at 30° C to an OD600 of 0.4, at which point the temperature was lowered to 18° C. Approximately one hour later overexpression was induced with 0.25 mM IPTG and left shaking overnight. Cultures expressing protein constructs containing the GR2 LBD were supplemented with 20 μM TA at the time of induction. After centrifugation, bacterial pellets were resuspended using a high salt buffer (25 mM Tris pH7.5, 1 M NaCl, 10 mM imidazole, 5% glycerol) supplemented with protease inhibitor tablets (Roche), 0.5 mM PMSF, and 2 mM beta-mercaptoethanol. For proteins containing the GR2 LBD 10 μM TA was present throughout the entire purification. Proteins were then purified using Ni-NTA affinity chromatography. At this point the purification was adjusted for the individual proteins.

GFP-tagged proteins were further purified using a Superdex 200 (Increase 10/300) column in 20 mM Hepes (pH 7.5), 250 mM NaCl, and 0.5 mM TCEP.

The DhL construct was cleaved using Ulp-1 enzyme. Protein was then diluted using 20 mM Tris (pH8.0) and 5% glycerol to reduce the salt concentration to less than 150 mM NaCl and loaded onto an assembly of an anion exchange column (HiTrap Q, GE) attached to the end of a cation exchange column (HiTrap SP, GE). After washing out unbound sample the anion exchange column was removed, and protein bound to the cation exchange column was eluted using a linear salt gradient. The protein was then concentrated and further purified using a Superdex 200 (Increase 10/300, GE) column in 20 mM Hepes (pH 7.5), 150 mM NaCl, 10 μM TA, and 0.5 mM TCEP.

GR constructs containing the GR NTD (full-length, NDh, and ND) were diluted using 20 mM Tris (pH8.0) and 5% glycerol to reduce the salt concentration to less than 100 mM NaCl and purified using an anion exchange column (HiTrap Q, GE). This was followed by purification over a Heparin column (HiTrap Heparin HP, GE), and finally proteins were purified using a Superdex 200 (Increase 10/300, GE) column in 20 mM Hepes (pH 7.5), 150 mM NaCl, 10 μM TA (for full-length GR only), and 0.5 mM TCEP.

### Fluorescent protein labeling

Proteins were prepared at 10 μM in 20 mM Hepes (pH 7.5), 250 mM NaCl, 0.5 mM TCEP and Cy3- or Cy5-maleimide (Fluoroprobes) were added at 30 μM final concentration. After incubation for 1 hour at room temperature proteins were purified using PD MiniTrap™ G-25 columns (GE Healthcare).

### Turbidity measurements

Turbidity of protein solutions was determined as the absorption at 340 nm using a Nanodrop OneC instrument (ThermoFisher). Protein samples were first prepared on ice at two times the desired final concentration in 20 mM Hepes (pH 7.5), 125 mM NaCl, 10 % glycerol, 0.5 mM TCEP, and 10 μM TA. Samples were then diluted 2-fold using the same buffer supplemented with 20% PEG8000. For measurements in the presence of DNA, protein samples were prepared as above at two times the desired final concentration and then diluted 2-fold with DNA at an equimolar concentration in buffer supplemented with 20% PEG8000. Turbidity was measured after a 10-minute incubation at room temperature.

### Droplet imaging

For fluorescent imaging Cy5- and Cy3-labelled proteins were mixed with non-labelled protein at a 1:50 ratio. Protein samples were then prepared on ice at two times the desired final concentration in 20 mM Hepes (pH 7.5), 125 mM NaCl, 10 % glycerol, 0.5 mM TCEP, and 10 μM TA. For measurements in the presence of DNA, GR was first incubated with DNA at equimolar concentrations for 5 minutes on ice before the addition of GFP-tagged co-regulators. Samples were then diluted 2-fold using the same buffer supplemented with 20% PEG8000. The resulting samples were then incubated at room temperature for 5 minutes and loaded onto a homemade chamber comprising a glass slide with a coverslip attached by two parallel strips of double-sided tape. After another 5 minutes incubation at room temperature droplets were imaged on the surface of the coverslip using a wide-field microscope (Olympus IX81 and Slidebook software; 100× magnification [UPIanFI, 1.30 NA oil]).

### Image analysis of droplets

To analyze phase separation imaging experiments, at least 5 images were collected and analyzed for each particular condition. Droplets were identified in *FIJI* as regions of interest using the “Analyze Particles” function in the Cy5-GR channel. Droplet areas and mean signal intensities for both channels were then determined and further analyzed in *R* using *RStudio.* Signal ratios were determined by dividing the mean GFP-signal by the mean Cy5 signal for each droplet. For normalization of the signal ratio data in the experiments using different DNAs, the signal ratios were normalized for each condition by dividing by the median signal ratio of the condition without DNA.

### Protein cross-linking

10μM NDh, 20μM GR2-LBD, and 10μM DNAs were mixed as indicated in buffer containing 20μM HEPES (pH 7.5), 125 mM NaCl, 0.5 mM TCEP, and 10μM TA. 0.5 mM BS3 cross-linker was added and the reaction was incubated at room temperature for 15 minutes before quenching with a final concentration of 100 mM Tris (pH 8.0). Samples were analyzed by SDS-PAGE.

### Reporter gene assays

Reporter gene assays were performed in U-2 OS human osteosarcoma cells, which were maintained and passaged in α-minimal essential medium (Life Technologies) supplemented with 10 % stripped fetal bovine serum (Invitrogen). Cells were transfected with 10 ng of hGR WT or K5xA mutant, 50 ng of SGK1, PLAU or TSLP firefly luciferase reporter(Hudson et al., 2018a), and 1 ng of Renilla luciferase reporter with FuGene HD (Promega) in OptiMEM (Invitrogen) according to the manufacturer’s protocol. Cells were treated with 100nM of TA or DMSO 24 hours after transfection in triplicate. Renilla and firefly luciferase activities were measured 24 hours after drug treatment using the DualGlo kit (Promega) by a BioTek Neo plate-reader (Winooski, VT).

## Author Contributions

F.F. and E.A.O. designed and conceived the study. F.F. performed all *in vitro* experiments and data analysis. X.L. performed reporter gene assays and data analysis. F.F. and E.A.O. wrote the manuscript. All authors reviewed the manuscript.

