## Supplementary_Information for "Glucocorticoid receptor condensates link DNA-dependent receptor dimerization and transcriptional transactivation"

Filipp Frank[1*], Xu Liu[1], Eric A. Ortlund[1*]
[1]Department of Biochemistry, Emory University School of Medicine, Atlanta, GA 30322, USA

*Corresponding author:
Filipp Frank,
,
Emory University,
Department of Biochemistry,
1510 Clifton Rd. NE, Atlanta, GA, 30322, USA.

Eric A. Ortlund,
,
Emory University,
Department of Biochemistry,
1510 Clifton Rd. NE, Atlanta, GA, 30322, USA.

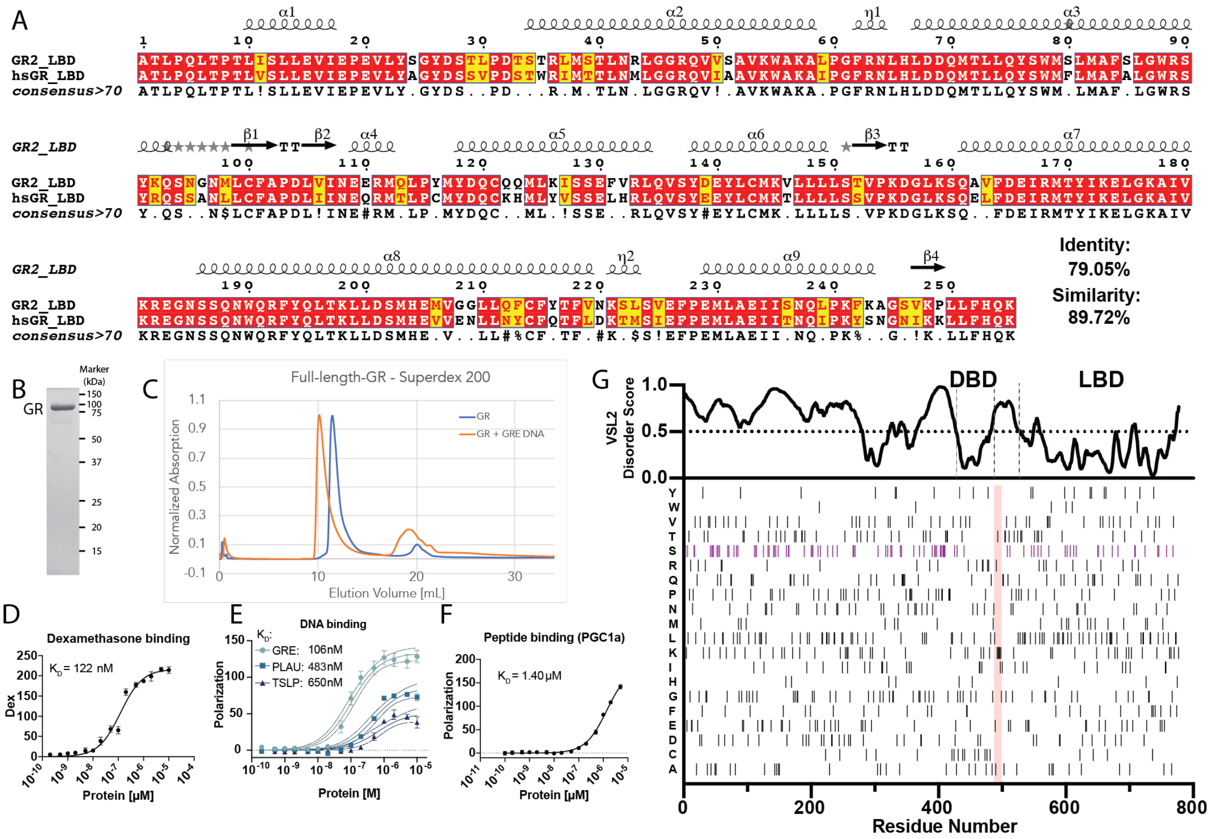


Figure S1: Recombinant full-length GR containing the ancestral GR2 LBD. (A) Sequence alignment between the human GR and ancestral GR2 LBD. (B) Coomassie blue stained SDS-PAGE gel showing purified full-length GR containing the GR2 LBD. (C) Superdex 200 size exclusion profiles for GR and a complex of GR with GRE DNA. (D-F) GR binding curves for interactions with Dexamethasone (D), different DNA elements (E), and co-regulator peptide (F). (G) Sequence analysis of GR showing that the N-terminal domain and the hinge region are intrinsically ordered. Bottom: Amino acid distribution along the GR sequence shows enrichment of serines in the N-terminal domain.

This figure is related to main text Figure 1.

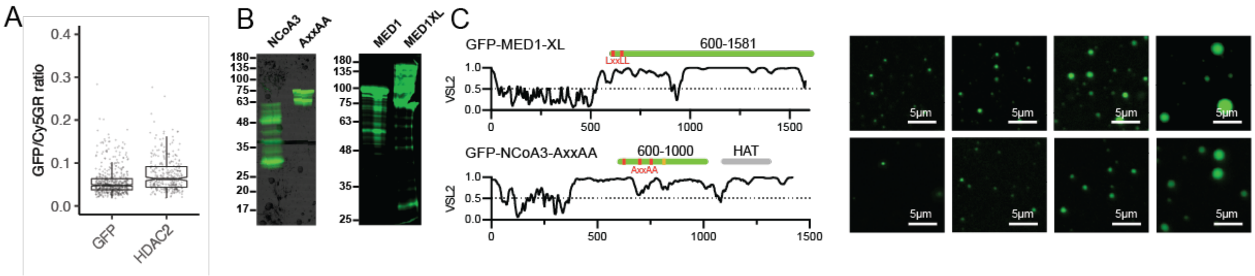

Figure S2: Phase separation behavior of GR co-regulator IDRs. (A) 10 µM GFP and GFP-HDAC2-IDR were mixed with 10µM GR and the recruitment of GFP and GFP-HDAC2-IDR into GR droplets was quantified by imaging. (B) Fluorescence images of SDS-PAGE gels with GFP-tagged IDRs as indicated. For droplet recruitment experiments equal amounts of proteins were added as determined by GFP-signal (absorption at 490 nm). (C) GFP-MED1XL and GFP-NCoA3-AxxAA constructs. LxxLL motifs are indicated in red. NCoA3 contains an additional LxxIL motif (shown in orange) which was also mutated to AxxAA. Images on the right show droplet formation of these constructs in droplet formation buffer (25 mM Hepes (pH7.5), 125 mM NaCl, 10% glycerol, 0.5mM TCEP, 10% PEG8000) at increasing concentrations (1µM, 3µM, 10µM, 30µM). This figure is related to main text Figure 2.

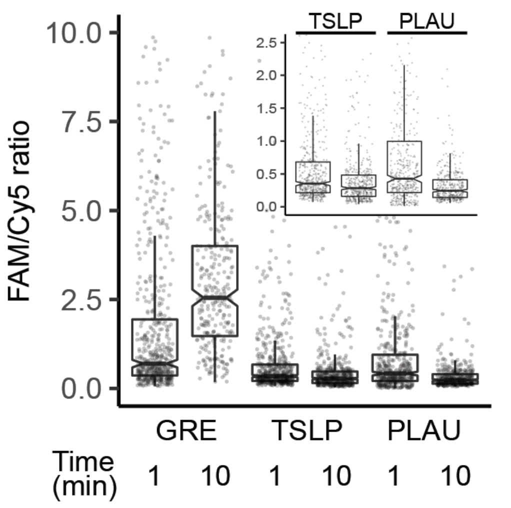


Figure S3: Quantification of the droplet mixing experiments shown in main text Figure 3 F and G. Droplets of GR bound to Cy5-labelled GRE DNA were mixed with droplets of GR bound to FAM-labelled GRE, TSLP, or PLAU DNAs and the distribution of fluorescence signals was followed over time. Shown is the ratio of FAM and Cy5 fluorescence signals of droplets identified in the Cy5-GRE channel measured right after mixing (1 minute) and 10 minutes later. When mixing FAM-GRE with Cy5-GRE the signal ratio increases over time because the droplets dynamically interchange with each other so that FAM-GRE is recruited into Cy5-GRE containing droplets (and vice versa). When mixing FAM-PLAU or FAM-TSLP with Cy5-GRE, however, the signal ratio decreases over time. This is due to the FAM-labelled repressive DNAs (TSLP and PLAU) being excluded from Cy5-GRE containing droplets, which results in diffuse FAM background signals.

This figure is related to main text Figure 3.

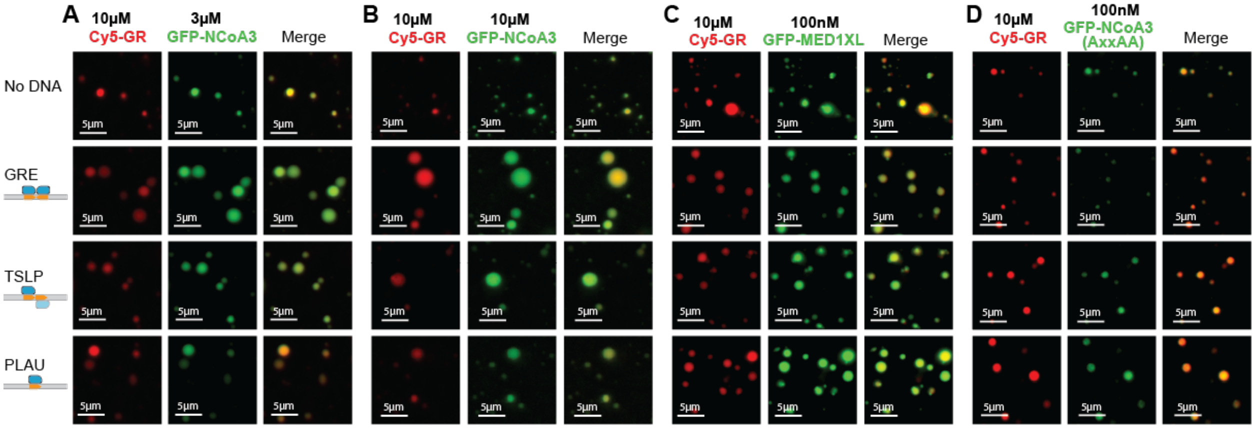


Figure S4: Representative images of GR droplets formed in the presence of GFP-tagged co-regulator IDRs as indicated. (A and B) Formation of GR droplets in the presence of high concentrations of NCoA3 does not induce the three-phase behavior seen for G9a and MED1. (C and D) Formation of GR droplets in the presence of GFP-MED1XL (C) and the GFP-NCoA3-AxxAA mutant constructs (D).

This figure is related to main text Figure 4.

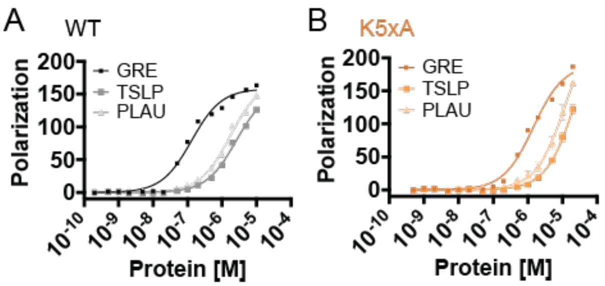


Figure S5: Characterization of the K5xA mutant. Shown are DNA binding curves of wild type DhL construct (A) and the corresponding K5xA mutant (B) with the indicated DNA elements.
This figure is related to main text Figure 5.
